# Rhythmic oscillations in the midbrain dopaminergic nuclei in mice

**DOI:** 10.1101/2022.12.24.521864

**Authors:** Virginie Oberto, Jumpei Matsumoto, Marco Pompili, Ralitsa Todorova, Francesco Papaleo, Hisao Nishijo, Laurent Venance, Marie Vandecasteele, Sidney Wiener

## Abstract

Dopamine release in the forebrain by midbrain ventral tegmental nucleus (VTA) and substantia nigra pars compacta (SNpc) neurons is implicated in reward processing, goal-directed learning, and decision-making. Rhythmic oscillations of neural excitability underlie coordination of network processing, and have been reported in these dopaminergic nuclei at several frequency bands. This paper provides a comparative characterization of several frequencies of oscillations of local field potential and single unit activity, highlighting some behavioral correlates. We recorded from optogenetically identified dopaminergic sites in mice training in operant olfactory and visual discrimination tasks. Rayleigh and Pairwise Phase Consistency (PPC) analyses revealed some VTA/SNc neurons phase-locked to each frequency range, with fast spiking interneurons (FSIs) prevalent at 1-2.5 Hz (slow) and 4 Hz bands, and dopaminergic neurons predominant in the theta band. More FSIs than dopaminergic neurons were phase-locked in the slow and 4 Hz bands during many task events. The highest incidence of phase-locking in neurons was in the slow and 4 Hz bands, and occurred during the delay between the operant choice and trial outcome (reward or punishment) signals. These data provide a basis for further examination of rhythmic coordination of activity of dopaminergic nuclei with other brain structures, and its impact for adaptive behavior.

## 1 Introduction

Dopamine is implicated in reward signaling, goal-directed learning, and decision-making, and related dysfunctions are linked with psychiatric disorders. The midbrain ventral tegmental nucleus (VTA) and substantia nigra pars compacta (SNpc) are principal sources of dopamine to the forebrain, and exert vital influences in the prefrontal-basal ganglia circuit via both neuromodulation and neurotransmission (Cerpa, et al., 2021; Chau, et al., 2018; Ott and Nieder, 2019; Williams, et al., 2002). These can have an impact on cognitive functions by affecting coherence of rhythmic oscillations of local field potentials, modulating the timing of neuronal discharges, and leading to synchronous activation of neurons as assemblies (e.g., Benchenane, et al., 2010). Furthermore, increased dopamine leads to changes in amplitude of oscillations in downstream structures (Berke, 2009). Such coordinated activity between different brain structures is associated with executive functions (Gray, et al., 1989; Varela, et al., 2001; Womelsdorf, 2007; Oberto, et al., 2021). In the case of the dopaminergic nuclei, such synchrony could be instrumental for the credit assignment problem: which synaptic strengths to reinforce or diminish following reward (or missed expected reward)? Rhythmic oscillations of field potentials and rhythmic activity of neurons have been reported in the VTA-prefrontal cortex-amygdala circuit, principally in the 4 Hz, theta and gamma bands. The 4 Hz rhythm there is associated with working memory (Fujisawa and Buszáki, 2011; Duvarci, et al., 2018) and modulates the 8 Hz theta and 50 Hz gamma rhythms. However, there are only rare reports concerning other prominent frequency bands present in VTA LFP: during sleep there is infraslow (Peters, et al., 2004) and delta (Gao, et al., 2007) rhythmicity while alpha band oscillations were once reported in behaving animals (Park and Moghaddam, 2017). None of this work has examined animals during instrumental tasks, however. There is some evidence for rhythms originating in the VTA (Peters, et al., 2004; Lowes, et al., 2021). Here, we identify and characterize a wide range of frequency bands operating in the brainstem dopaminergic nuclei through analyses of LFPs and neuronal phase-locking.

We find that multiple rhythms are prominent in the VTA/SNc in mice acquiring and performing a goal-directed task (specifically selected to elicit dopaminergic activity). These rhythms are observed in LFPs, and their role in local processing (as opposed to volume conduction) is evidenced by detection of phase-locking of spike trains of neurons at optogenetically confirmed recording sites. These rhythms could orchestrate interactions between VTA and an extended network of cortical and subcortical structures including striatum and prefrontal cortex (PFC).

## 2 Materials and Methods

### 2.1 Animals

All experiments were performed in accordance with EU guidelines (directive 86/609/EEC). Subjects were transgenic adult (2-9 months) male mice expressing a channelrhodopsin2-yellow fluorescent protein fusion protein (ChR2 (H134R)-eYFP). These mice were obtained by mating mice expressing the CRE recombinase under the control of the dopamine transporter (DAT-IRES-CRE mice, Stock 006660, Jackson Laboratory, ME, USA) with mice bearing a CRE-dependent ChR2(H134R)-eYFP gene (Ai32 mice, Stock 012569, Jackson Laboratory). Animals were housed in temperature-controlled rooms with standard 12-hour light/dark cycles and food and water were available *ad libitum*. Each workday, animals were handled to habituate to human contact, and weighed. These mice were obtained by mating mice expressing the CRE recombinase under the control of the dopamine transporter (DAT-IRES-CRE mice, Stock 006660, Jackson Laboratory, ME, USA) with mice bearing a CRE-dependent ChR2(H134R)-eYFP gene (Ai32 mice, Stock 012569, Jackson Laboratory). During pre-training and experimental periods, food intake, including pellets provided in the experimental apparatus, was restricted to a maximum of 3 g/day. Water remained ad lib. Supplemental food was provided if weight fell below 85% of the normal weight, taking into account the animal’s age.

### 2.2 Recording and stimulation

In order to compare activity along the mediolateral axis of the dopaminergic nuclei, an array of 4 recording probes was constructed attached to 2 optical stimulation probes, with each serving the medial or lateral pair of recording probes. Twisted bundles of 8 formvar coated 12 micron diameter nichrome wires (“octrodes”) were inserted in polyimide tubes. The wires were gold-plated to an impedance of about 350 kohm. The main body of the headstage is composed of a resin (MadeSolid, Oakland, USA) molded in a 3D printer. The drives were adapted from the “lambda drive” of Headley, et al (2015). Two optic fibers were inserted in the headstage, which positioned the medial two at a depth of 3.9 mm above VTA, and the third and fourth at 3.7 and 3.5 mm abouve SNc, with the tips facing rostrally. The entire assembly is shielded from electrical noise with a fine mesh spot-glued with dental cement. This was connected to ground, serving as a miniature Faraday cage. Recordings were made with an Ampliplex system (sampling rate: 20 kHz; cutoff frequencies of the analog low-pass and high-pass filters were 0.3 and 10 kHz, respectively). After surgical implantation, electrodes were advanced as often as twice daily until neurons with responses to optical stimulation appeared. Electrodes were further advanced when discriminable units were no longer present.

### 2.3 Surgery

Anesthesia was induced with a mixture of ketamine (66 mg/kg) and xylazine (13 mg/kg) and sustained with isoflurane (0.5 – 1.0 %). Mice were placed in a stereotaxic device (David Kopf Instruments) and maintained at 37° C. The scalp was exposed and cleaned with hydrogen peroxide solution. Miniature jeweler’s screws were screwed into trephines and attached with dental cement. The electrode assembly was implanted in the configuration described above into the left VTA and SN. The stainless-steel screws implanted on the left and right cerebellum as ground and reference electrodes, respectively.

### 2.4 Behavioral task

The ‘Operon’ system developed by Scheggia et al (2014) was adapted as an experimental chamber permitting the mice to perform olfactory and visual discrimination tasks (Figure 6A). Each of the short sides of the rectangular (160L x 136W x 160H mm) plexiglass chamber has two arrays of LEDs and two ports used for both odor sampling and nose pokes. An olfactometer presented carvone or its enantiomer, limonene, at the respective odor ports in a pseudo-random sequence. All six white LEDs were either lit or off above the respective odor ports, again in pseudo-random sequence. Between the ports and array is a central feeder port with an overhead lamp which is lit for 7 s during the delay after error trials. A reward pellet (5TUL 14mg, TestDiet) was delivered by a dispenser (ENV-203-14P, Med Associates). A plexiglass barrier divided the two sides of the chamber and was slid up from below via an automated motor assembly. At the beginning of each trial, the barrier was lowered. Once the mouse crossed a photodetector on its way to the other half of the chamber, one LED array was lit and two odors were released from the respective odor ports on that side. This photodetector crossing is called the “Cue” event. Once the mouse cleared these photodetectors, the barrier was slid shut.

Nosepokes to the port on the side of the (preselected) odor or visual cue crossed a photodetector within the port that triggered food pellet release, or an error signal, at a variable latency ranging from 0.5 to 1.0 s. The position of the rewarded cue was pseudorandomly varied on successive trials so that the same odor or visual cue were not presented more than twice at the left or the right nosepoke port. Furthermore, the same trajectory orientation across the chamber was not rewarded on more than 2 successive trials. The task control system permitted programming the cue contingencies and reward delivery, outputting time stamps. Erroneous choices triggered illumination of an overhead light during a timeout period of 5 or 8 s while the barrier remained up.

Animals were first pre-trained to shuttle between two boxes for reward. In the first stage of training, a nose poke into either port triggered a pellet delivery. Then animals were pre-trained for simple visual discrimination task (light-on vs light off). While the rewarded side was varied pseudo-randomly, in cases of spatial persistence, the other side was rewarded more. After implantation, training was resumed along with recordings.

Mice were provided water ad libitum, but mildly food deprived to maintain them above 85% their normal weight, as controlled with daily weighing. The behavioral apparatus was turned on and the olfactometer outputs were confirmed to present odors discriminable by the experimenter. The headstage was then plugged in, and the mouse was placed into one compartment of the conditioning chamber. The room lights were dimmed and the recording system was turned on and the behavioral task was started. In the early sessions, the animals were trained in the individual tasks. Once they attained proficiency in olfactory and visual discriminations, reaching criterion (8 correct choices out of 10 consecutive trials) performance in one task triggered the system to switch to the other task. The mice could then cycle between the two tasks. When the mouse stopped performing trials for more than 120 s, the behavioral session was ended. The headstage plug was changed, optic fibers were connected, and the animal was placed in a large plastic beaker. Using a 475 nm-laser light source (DPSSL BL473T3-100FL, Shanghai Laser and Optics Century), trains of 10 light pulses (pulse duration: 10 msec; light power: 10 mW max; frequency: 5 Hz) were given for 20 times with 10 sec inter train intervals under control of an Arduino-based system. The weight of the rewards earned was calculated, and the required amount of chow for the daily feeding was calculated and provided.

### 2.5 Data analyses

#### 2.5.1 Single unit discrimination

Offline spike sorting was carried out with KiloSort (Pachitariu et al., 2016) followed by manual curation using Klusters (L. Hazan, http://neurosuite.sourceforge.net).

To control for the risk that neurons stably recorded across days would be counted more than once, we compared electrophysiological features of pairs of neurons recorded from the same octrode on consecutive days. These features included spike waveform shape, spike auto-correlogram profiles, and mean firing rate with the method of Fraser and Schwartz (2012). To quantify the waveform similarity, the peak value of the cross-correlogram between the pairs of samples of average waveform shapes of the units was calculated for each channel of the octrode. The resulting coefficients of different channels were then averaged. The waveform similarity score was calculated as Fisher transform of these averaged coefficients. To quantify the auto-correlogram similarity, the auto-correlograms were calculated from 0 to 100 ms with 5 ms binning. Pearson correlation coefficients between the auto-correlograms of a pair of units were calculated and Fisher transformed. The resulting value is the auto-correlogram similarity score. The mean firing rate similarity score was calculated as the difference between the logs of the mean firing rates in the two recordings. The three similarity scores were combined with a quadratic classifier that computes an optimal decision boundary between same and different neuron pairs under the assumption that the underlying data can be modeled as a mixture of multivariate Gaussians. Since the distribution of different neuron pairs can be obtained using the data obtained from different octrodes, we used partially supervised expectation-maximization to fit a mixture of Gaussian models utilizing this prior information. This was used to estimate the distribution of same-neuron pairs (red dots and curves in Supp. Fig. 2). Specifically, the decision boundary of the classifier is calibrated to produce a 1% error rate in the known different-neuron distribution, and pairs with values lying inside the boundary were identified as same-neuron pairs. Then, different-neuron pairs recorded from same octrodes were estimated. To be conservative, those different-neuron pairs were estimated after the decision boundary of the classifier is re-calibrated to produce a 1% error rate in the same-neuron pair distribution estimated above (Supp. Fig. 2). For the data analysis, we used only a single recording day from each series of recording from a potentially same neuron (series of recordings connected with pairs represented by magenta (unknown pair) and red (same-neuron pair) dots in Supp. Fig. 2). Pairs not assigned as same or different were considered as ‘unclassified’.

#### 2.5.2 Neuron categories

To identify neurons as dopaminergic, we used the Stimulus-Associated spike Latency Test (SALT; Kvitsiani et al., 2013; Eshel et al., 2013). The test determines whether light pulses significantly changed a neuron’s spike timing by comparing the distribution of first spike latencies relative to the light pulse, assessed in a 10-msec window after light-stimulation, to 10-msec epochs in the baseline period (−150 to 0 msec from the onset of light-stimulation; see Kvitsiani et al., 2013 for details). A significance level of p<0.01 was selected for this.

All neurons recorded from an octrode with at least one SALT+ response were considered to be in a dopaminergic nucleus. If the octrode was not advanced, neurons on the day before and the day after were also counted as in VTA or SNc, even if no SALT+ responses were recorded on those days. Similarly, if the octrode had not been advanced, and SALT+ responses had been recorded on non-consecutive days, intervening days with no SALT+ responses were still considered as in a dopaminergic nucleus. Also, when electrodes had been advanced, all recordings in days between those with SALT+ recordings were also considered to be in VTA/SNc. Neurons with SALT+ responses are considered dopaminergic. Neurons in a dopaminergic nucleus not demonstrating a significant SALT+ response are labelled SALT-, although this negative result does not conclusively show that the neuron is not dopaminergic. All neurons qualifying as in dopaminergic nuclei were categorized with clustering according to the following parameters: spike width, mean firing rate. The criterion for fast spiking interneurons (FSI) was firing rate > 15 Hz and spike width < 1.5 ms (cf., Ungless and Grace, 2012). No FSI’s were SALT+. All other SALT-neurons in dopaminergic nuclei are referred to as “Other”.

#### 2.5.3 LFP recordings and phase-locking in neurons

LFPs were derived from wideband signals that had been down-sampled to 1250 Hz on all channels. Power spectra were calculated with wavelet methods. LFP was taken from the median voltage of all active channels of an octrode containing at least one channel with SALT+ responses. Spikes were removed from the LFP signal according to the method of https://www.fieldtriptoolbox.org/tutorial/spikefield/#analyzing-spikes-and-lfps-recorded-from-the-same-electrode. We quantified phase consistency of spikes relative to the LFP band with both the Rayleigh test of circular uniformity and the unbiased pairwise phase consistency (PPC, Vinck et al 2010). With the Rayleigh test, a neuron was considered as phase-locked if p < 0.05. In the second method, PPC threshold was determined from a shuffling analysis: For each neuron and each perievent window, interspike intervals were shuffled for each trial. The actual values were considered significant if they exceeded the 95th percentile of this distribution of PPCs calculated for the shuffled data. Response profiles were similar for Rayleigh and PPC tests, and most results are given for the latter.

To control for potential influence of different numbers of spikes in samples on PPC results (e.g., due to differences in firing rates among neurons, or during the respective task events), PPC values and significance thresholds were calculated on equal sample sizes by down-sampling the data. PPC was calculated on 50 spikes randomly selected from the task event period or, for analyses of the entire session 250 spikes. This procedure was repeated 1000 times, and resultant PPC values are averaged to calculate the subsampled PPC value. The threshold for testing the significance of the PPC value (p < 0.05) was determined by bootstrapping the PPC results using shuffled spike data (Thorn and Graybiel, 2014). Median PPC’s were calculated for neurons tested that had 50 or more spikes for the period examined.

#### 2.5.4 Behavioral analyses

Data were analyzed for [-500 ms, 500 ms] periods around three trial events. “Cue” is when the mouse first crossed the photobeam while entering the next chamber, triggering cue onset. “Choice” is the time of the nosepoke response, as detected by the photo-detectors in the ports. “Outcome” is either the instant the dispenser released the reward pellet (which made a salient sound), or the onset of the punishment period. For PPC and Rayleigh analyses, eight trial periods are examined and compared: pre-cue [cue onset-0.5 s, cue onset], post-cue [cue onset, cue onset+0.5 s], and corresponding periods for the choice onset, reward onset and punishment onset events.

#### 2.5.5 Histology

Once stable recordings were no longer possible, marking lesions were made with 10 s of 30 µA cathodal current. After waiting at least 90 min, mice were then killed with a lethal intraperitoneal injection of sodium pentobarbital, and perfused intra-ventricularly with phosphate buffered saline solution (PBS), followed by 10% phosphate buffered formalin. The brain was removed, post-fixed overnight, and placed in phosphate buffered 30% sucrose solution for 2-3 days. Frozen sections were cut at 80 µm, and permeabilized in 0.2% Triton in PBS for 1 h at room temperature. They were then treated with 3% bovine serum albumen (BSA) and 0.2% Triton in PBS for 1 h with gentle agitation at room temperature to block non-specific binding. Sections were then rinsed for 5 min in PBS at room temperature with gentle agitation. Then the sections were left overnight with gentle agitation at 4° C in a solution of the first antibody, mouse monoclonal anti-TH MAB318 (1:500), 0.067% Triton, and 1% BSA in PBS. After three rinses for five minutes in PBS, the sections were treated with a second antibody (1/200, anti-mouse, green), Nissl-red (1:250), and 0.067% Triton in PBS for two hours at room temperature. Then sections were rinsed 3 times for five minutes in PBS, and mounted with Fluoromount®. Sections were examined with fluorescence microscopy.

## 3 Results

### 3.1 Determining the principal frequency bands of LFP oscillations and neuronal phase-locking in VTA/SNc

In recordings from microelectrodes confirmed optogenetically to be in the midbrain dopaminergic nuclei (Supp. Figure 1), several rhythms were readily apparent even in raw unfiltered LFP traces (Figure 1). To verify this, spectra were analyzed to determine the principal frequency bands most prominent in these nuclei, without making any a priori assumptions of their ranges (Figure 2). Peak frequencies were identified as inflection points in this spectrum, visualized as local minima in the second derivatives of these traces. Their frequencies were 0.7, 4, 10, 22, and 40 Hz (Figure 2). To verify this, a FOOOF analysis (Donoghue, et al., 2020) was performed, and curve-fitting yielded bands centered at 4, 8, 41, and 50 Hz (not shown). These results derive exclusively from LFP traces, which carry the risk of contamination with volume-conducted signals. Thus, single unit recordings were analyzed for evidence of rhythmic activity intrinsic to the dopaminergic nuclei.

**Figure 1.** Identification of principal bands in recordings from the same octrode in VTA in a behaving mouse. Top row) Unfiltered traces. Cue, choice and reward refer to the three task events. Examples of the oscillations in the frequency bands are circled in the top trace, and in their respective filtered traces below. All y-axes are voltage in arbitrary units (a.u.).

**Figure 2.** Detection of the dominant LFP frequencies in the midbrain dopaminergic nuclei. Top) Mean (black trace) of recordings from all octrode recordings in optogenetically identified dopaminergic nuclei. Gray traces represent SEM’s. 1/f correction was made. Middle and bottom) First and second derivatives of top trace. Centers of the respective frequency bands (indicated at the top of the Figure) correspond to local minima in the bottom plot.

We recorded in mice training and performing sensory discrimination tasks (described further below). The neurons reported here were recorded from octrodes with at least one positive dopaminergic response (SALT+), or on days between days with dopaminergic responses, and when the electrodes had not be advanced. There were 171 dopaminergic neurons (DA), 113 FSI neurons, and 375 Other neurons (a total of 659 neurons). Neurons were excluded when there was evidence that they had been repeatedly recorded on successive days (see Material and Methods). After this, there remained 85 dopaminergic neurons (DA, SALT+), 41 fast spiking inter neurons (FSI), and 254 SALT-non-FSI (Other). Note that the latter group could include dopaminergic neurons whose optogenetic responses were not significant (because of possible technical issues such as insufficient illumination or expression of opsins).

Rayleigh and pairwise phase consistency (PPC; Vinck, et al., 2010) analyses assessed neuronal activity phase-locked to LFP oscillations. Frequency bands with narrow windows were examined, and the sliding windows passed successively through the range from 0.1 to 100 Hz. The incidences of significant results were tallied, and several frequencies appeared repeatedly in both tests (Figure 3). These were primarily centered around bands similar to those previously reported elsewhere in the brain: 1-2.5 (slow), 2.5-6.5 (4 Hz), 6.5-13 (theta), 13-30 (alpha), 25-55 (low gamma), and 40-75 Hz (high gamma). These bands are also largely consistent with the LFP spectrum analyses, and thus are selected for further analyses below.

**Figure 3.** Determining principal frequency bands from oscillatory entrainment in neurons (all cell types pooled together) in the brainstem dopaminergic nuclei. Top row) Significant responses in Rayleigh test, PPC or both for each neuron (rows) at various frequencies (tested with a sliding scale). The color scale is normalized for each neuron. Middle row) Cumulative summary of counts of significant neurons from above. Bottom) Comparison of results from the middle row.

### 3.2 Characterization of neuron phase-locking to rhythms

Figure 4 shows examples of neurons phase-locked to the respective frequency bands. For the example in the slow band, the spike triggered LFP averages (STA) completed a full cycle 0.6-0.8 s after the spike time, corresponding to frequencies of 0.6-0.8 Hz. STA cycle lengths for the other frequency bands are highlighted by arrows above the plots, and these correspond to the respective frequency bands. For 4 Hz, the auto-correlogram of the example on the left in Figure 4 shows a prominent peak corresponding to this frequency band. For the theta band, both examples have auto-correlograms and spike-triggered average with delays on the order of 100 ms, corresponding to 10 Hz.

**Figure 4.** Examples of neurons in the midbrain dopaminergic nuclei with phase-locking to the principal frequency bands. Data were recorded during performance of the T-maze task and are from periods from 5 s prior to cue onset until 5 s after the trial outcome (reward or punishment) over all session trials. Two examples (left and right columns) are shown for each frequency band. Top and 4^th^ row) Average waveform from one octrode channel (scale bar is 1 ms). 2^nd^ and 5^th^ rows) Spike auto-correlograms show peaks, bumps or shoulders at delays corresponding to the respective frequency bands (highlighted with vertical bars and arrows). 3^rd^ and 6^th^ rows) Phase-locking of spike activity to the respective frequency bands. Cosine wave above is schematic. 4^th^ and 7^th^ rows) Spike-triggered averages (STA) of the LFPs filtered to the respective bands (arbitrary units). Arrows and vertical bars above STA’s show intervals corresponding to the respective frequency bands. For 1.0-2.5 Hz, both neurons were recorded from octrodes with at least one optically identified dopaminergic neuron but did not qualify as dopaminergic or FSI (thus referred to as unclassified); 2.5-6.5 Hz are unclassified (left) and DA (right); 6.5 −13 Hz are unclassified (left) and FSI (right); 13-25 Hz and 25-55 Hz are both DA; and 40-75 Hz are DA (left) and unclassified (right).

The greatest numbers of phase-locked neurons were in the slow and 4 Hz bands for the FSIs, but these did not have a significantly greater incidence than in the other bands. However, for DA neurons, there was a greater incidence of phase-locking in the theta band than slow or beta bands (binomial test, p<0.05 with Bonferroni correction with Bonferroni correction).

We next tested whether neurons were phase-locked to multiple bands in PPC analyses (e.g., two adjacent frequency intervals). Supp. Fig. 3 shows that low proportions of neurons were phase-locked to more than one band, indicating that the selected frequency bands are discrete and independent.

PPC analyses of whole session data examined the incidence of phase-locking of DA and FSI to the respective bands. This was significantly greater in FSI than DA neurons, in the slow and 4 Hz bands (Figure 5A; binomial test p<0.05 with Bonferroni correction). While in the theta band, a greater number of DA neurons were phase-locked than FSI’s, this was not significant. This issue will be further addressed below.

**Figure 5.** A) Incidence of PPC phaselocking in DA and FSI over the entire task. B) Distribution of preferred phases of DA (black) and FSI (white). Circular means in radians [DA, FSI] are: slow [n.s., 3.1], 4 Hz [0.8, 2.2], theta [1.0, 2.2], beta (not shown) [n.s., n.s.], low gamma [2.4, 2.5], high gamma [2.9, 1.8]. Overlapping zones are in gray. *-p<0.05

We then examined the preferred phases of FSI and DA for the respective bands for all neurons (excluding repeat recordings; Fig. 5B). Rayleigh tests revealed significant (p<0.05 with Bonferroni correction) preferred phases for both neuron types for all bands except for the beta band, and DA in the slow band. The preferred phases of FSI and DA were significantly different at 4 Hz, theta and high gamma (Kuiper test, p<0.05) but not low gamma (which had a relatively low sample).

### 3.3 Behavioral correlates of LFP amplitude and phase-locking of neurons in VTA/SNc

Next we examined the evolution LFP amplitude and incidence of phase-locking during acquisition of visual and odor discrimination tasks (described in section 2.4; see Figure 6A). Since the respective phases of task execution would call upon different cognitive processes, requiring synchronization of various circuits, analyses compare event-related oscillations and phase-locking of neurons. As the mice performed a trial, they first entered the opposite compartment, crossing a photo-detector, which triggered onset of visual or olfactory cues, or both. This is the “cue” event, and the 500 ms periods prior and after this involved approach behavior (“pre-cue”) and cue detection and analysis (“post-cue”). The next event was the nose-poke response to the left or right odor ports, again crossing a photo-detector. The 500 ms events prior and after corresponded to decision (“pre-choice”) and anticipation of reward or punishment (“post-choice”). There was a variable delay between the nose-poke and either the click of the food dispenser for rewarded trials, or the onset of the punishment signals. These delimited the 500 ms “pre-outcome” and “post-outcome” periods.

**Figure 6.** A) Schematic of the behavioral chamber (adapted from Scheggia, et al., 2014). B) LFP during task periods for rewarded trials only (corrected for 1/f). There are discontinuities because the time between successive events varied among trials. Left) Mean spectrograms from all data synchronized with the 3 principal task events. Right) Breakdown for the respective bands. Data here are only from rewarded trials. C) Incidence of PPC significant neurons in the respective task periods (DA and FSI pooled together). *-p<0.05; **-p<0.01; *** - p<0.005.

#### 3.3.1 The slow band

In the slow band, LFP amplitude was maximal during the cue presentation period and lowest during the outcome period (Figure 6B). A one-way ANOVA of PPC values of FSI’s across trial periods was significant (Kruskal-Wallis, p=0.029), with the values in the pre-reward period greater than the pre-cue period (binomial test; p<0.05 with Bonferroni correction, reward and punishment outcomes were analyzed separately here). Thus, even though LFP amplitude is relatively lower during this trial period, cells still can maintain regular phase relationships then. In all neurons pooled together, in the slow band there were significantly more neurons phase-locked in the pre-outcome period than the pre-cue period (Figure 6C). Detailed phase-locking results for this and other bands are in Supp. Fig. 4).

#### 3.3.2 The 4 Hz band

In the 4 Hz band, LFP amplitude was maximal during the cue period and lowest during the outcome period (Figure 6B). PPC values of FSIs are significantly different among task event periods (Kruskal-Wallis, p<0.001), with the post-choice period significantly greater than the pre-cue period (binomial test; p<0.05 with Bonferroni correction). A greater proportion of neurons phase-locked during the post-choice period than several other periods (Figure 5D; both neurons types pooled together). This tendency is apparent, but not significant, when DA and FSI are examined separately (Supp. Fig. 4).

#### 3.3.3 The theta band

Theta LFP amplitude peaks prior to the nose-poke choice and has local minima pre-cue and post-choice. Pairwise phase consistency (PPC) is significantly different between task periods for FSIs (Kruskal-Wallis, p=0.004; not shown).

#### 3.3.4 The beta and gamma bands

Beta LFP amplitude has local maxima at the nose-poke choice and again 300 ms after, and has local minima pre-cue and post-choice. Low and high gamma remain relatively stable, with a peak in low gamma 400 ms after the choice, figuring prominently in the spectrogram (Figure 6B).

There were no significant differences between incidence of significant phase-locking across the task events or between DA vs FSI for beta or gamma bands.

Note that the samples reported here (e.g., FSI vs. DA, frequency band comparisons) are conservative, since neurons determined to be repeatedly recorded in successive sessions were excluded. Also, differences in firing rates are taken into account for PPC, where we subsampled spikes (see Methods). While Rayleigh results are not reported above, the patterns of distributions of incidences of neurons significant for the Rayleigh test appear to resemble the PPC results (Supp. Fig. 4).

### 3.4 Phase-locking dependence upon task performance level

Since the task involved acquisition of new cue-reward associations, for which the dopaminergic nuclei are strongly implicated, we assessed if phase-locking changed as a function of performance level of the animals. We computed the proportion of phase-locked neurons in sessions where the animal reached criterion performance in at least one of the tasks, triggering the rule to be changed (n=96). These were compared to other sessions where mice did not achieve criterion performance (n=33). Significant differences were rare, and were not concentrated on any particular frequency band or trial event.

Other tests showed no differences in incidences of phase-locking of cells or of PPC values in comparisons of rewarded vs punished trials or between cues presented (visual vs odor vs both; p<0.05 with Bonferroni correction; data not shown).

## 4 Discussion

LFPs in the mouse dopaminergic nuclei showed prominent oscillations in several frequency bands. These were not simply volume conducted signals originating elsewhere since neurons recorded in optogenetically identified dopaminergic sites were also modulated by these rhythms. While 4 Hz, theta and low gamma have been documented, this is the first report, to our knowledge, of slow and beta oscillations in the dopaminergic nuclei of behaving animals.

Furthermore, behavioral correlates are reported concerning the amplitude of these oscillations, and phase-locking in neurons. Below these are discussed in relation to the published literature for each frequency band.

The distinct nature of these bands is supported by the minor incidence of neurons with phase-locking to more than one band. This tendency for exclusivity is consistent with the low incidences of neurons with phase-locking in the respective bands. If the respective frequency bands correspond to communication channels for different functional networks, this would suggest that these neurons have functional specificities.

Results were similar between Rayleigh tests and PPC, although more cells had significant responses with the former, especially when spike counts were high. This is likely because the Rayleigh test determines if there is a deviation from equal firing at all phases of the oscillation, while PPC determines whether there are common phases among spikes. Since we were interested in phase-locking during brief intervals related to the respective task events, and this reduced the number of spikes that could be analyzed, we selected PPC (which can yield results for as few as 30 spikes; Womelsdorf, et al., 2014) for most of these analyses. The Rayleigh test requires a minimal number of spikes for each bin of phase ranges to avoid false positives.

### 4.1 The slow rhythm

The frequency bands here were determined with several approaches based upon the actual data. This led to distinguishing separate bands for slow and 4 Hz. In some work, the delta band is considered to range from 1 to 5 Hz, and this could lead to confounds. For this reason, we did not use the term “delta” to describe the 1 to 2.5 Hz band here.

Overall, reports in the literature of slow oscillations during waking behavior are relatively sparse. But, the incidence of phase-locking to the slow rhythm here tended to be greater than the other bands, although this difference was not significant.

There are reports of cortical delta rhythms in awake rodents during stationary immobility in the absence of experimental sensory stimulation (Crochet and Petersen, 2006; Schultheiss, et al., 2020). However, these results are not directly comparable here since the range of their delta frequency band included 4 Hz. With intracranial recordings, Halgren et al (2018) found 2 Hz oscillations that were generated in the superficial layers of the cerebral cortex of awake epilepsy patients. These were coupled to higher frequency oscillations in other layers. We found slow rhythms in hippocampus and prefrontal cortex of rats performing in a T-maze, and noradrenergic locus coeruleus neurons were phase-locked to these oscillations (Xiang, et al., In chloral hydrate anesthetized rats Peters, et al. (2004) and Gao, et al. (2007) observed infraslow (0.65 and 0.86 Hz respectively) oscillations in DAergic neurons. They considered this as an experimental model for delta rhythms during sleep. Intracellular PFC UP states (Peters, et al. 2004) and PFC LFP negative deflections (Gao, et al., 2007) were synchronous with negative deflections of the VTA LFP at this frequency range. The VTA neurons fired in phase with high, but not low amplitude prefrontal cortical oscillations at this frequency. Non-DA neurons were also modulated, but were almost pi radians (180°) out phase from the DA neurons (Gao, et al. 2007). In contrast, in our awake behaving mice, the FSI and SALT-Other neurons had a phase difference of only 0.8 and 1.2 rad respectively from DAergic neurons. The VTA LFP also highly resembled that of PFC. TTX infusion in PFC suppressed the slow oscillations in VTA neurons (Gao, et al., 2007). Interestingly, lidocaine infusion in VTA (but not substantia nigra) suppressed the UP-DOWN state oscillations in PFC.

More FSIs than DA were phase-locked to the slow and 4 Hz rhythms. Interneurons are known to be instrumental in the generation of theta and gamma rhythms. Our experimental design was not appropriate for testing whether the rhythms are generated locally. While the higher firing rates of interneurons relative to principal neurons could render the PPC and Rayleigh tests more sensitive, this result remained robust since we down-sampled all spikes for PPC.

More neurons were phase-locked to the slow rhythm during the pre-outcome period than the pre-cue period. This could be related to synchronizing dopaminergic structures with associated structure during the period when reward is being anticipated.

Van der Velden, et al. (2020) optogenetically stimulated lateral VTA in mouse brain slices. Varying the frequency of stimulation, they found that individual neurons had optimal responses (“resonance”) at several values between 0.5 to 5 Hz. This corresponds to the slow band observed here, extending to the 4 Hz band as well. The authors suggest these variations in preferred infraslow and slow frequencies among individual neurons could support temporal processing of inputs and coding for outputs.

### 4.2 The 4 Hz rhythm

Fujisawa and Buzsáki (2011) showed that the power of 4 Hz oscillation (filtered from 2-5 Hz) in rat PFC and VTA, and its coherence between these two structures, are stronger during the choice phase of a working memory task. They found that 46% of putative dopaminergic and 38% of putative GABAergic VTA neurons were significantly phase-locked to the 4 Hz oscillation.

Similarly, 4 Hz rhythmicity (with 3-6 Hz filters) occurs in mice during choice periods in another working memory task (Duvarci, et al., 2018). 75% of DAergic, and 78% of GABAergic VTA cells were modulated by VTA 4 Hz. (In these papers, neuron types were identified according to responses to systemic apomorphine administration, baseline firing rates and spike width.) In contrast, here, the incidence of neurons with significant 4 Hz PPC was significantly elevated during the post-choice period (during the delay prior to reward or punishment). Discrepancies with the cited results could be related to our task lacking an explicit trial-by-trial working memory component. Another difference is that here, twice as many FSI’s than DAergic neurons were modulated by 4 Hz oscillations. This difference could be related to different methods used to identify the two types of neurons, the different filter range used, or sampling properties of the different types of electrodes used. Also, these other papers used the Rayleigh test to detect phase-locking, while we reported PPC results, which tended to have fewer significant FSI’s in this band (Supp. Fig. 4).

In Duvarci et al. (2018), the median preferred phase of DAergic neurons was about −2 rad, while GABAergic neurons were about −0.75 rad. In Fujisawa and Buzsaki (2011), relative to PFC LFP, DAergic neurons had a median phase of about −0.5 rad and GABAergic neurons had a median phase of about −2.6 rad. Here the mean preferred phase of DA neurons in the corresponding frequency band was 0.8 rad, but for FSI’s the value was 2.2 rad.

Lowe et al (2021) found increased amplitude of 4 Hz amplitude in VTA and nucleus accumbens (NAc) LFPs during stressful restraint in mice (which also reduces reward seeking). Furthermore, multi-unit activity in VTA was phase-locked to NAc 4 Hz. Interestingly, silencing VTA GABAergic neurons with muscimol suppressed NAc LFP 4 Hz in stressed animals.

One potential confound for 4 Hz LFP in mice is the correlation with breathing signals sent from olfactory bulb through piriform cortex and PFC (e.g., Bagur, et al., 2021). Rhythmic firing in the DAergic neurons here provides evidence that this is not simply a volume-conducted signal. Thus, the DA nuclei are part of the widespread network of brain structures coordinated by the 4 Hz rhythm (Tort, et al., 2018).

### 4.3 The theta rhythm

In behaving rodents, hippocampal theta is typically reported at 7-8 Hz. Here, the spectral data revealed an inflection point at 10 Hz, and thus the range studied was 8 to 13 Hz, and this helped to avoid overlap with the band centered on 4 Hz. Fujisawa and Buzsáki (2011) filtered for theta in their hippocampal LFP recordings at 7-11 Hz, but did not report VTA LFPs in this frequency range. However, they did observe VTA neurons modulated both by VTA 4 Hz (as replicated here) and by hippocampal theta. Future work should test for potential relations between hippocampal theta and VTA 10 Hz oscillations. Note that Park and Moghaddam (2017) filtered rat VTA LFPs for theta between 5-15 Hz, and found peaks at 10-12 Hz, consistent with the present work in mice. Fujisawa and Buszáki (2011) found significant phase locking to the hippocampal theta rhythm in 44% of putative dopaminergic neurons and 39% of putative GABAergic cells. In the Park and Moghaddam (2017) study, 45% of DAergic neurons were modulated by VTA 5-15 Hz, while only 23% of non-DAergic neurons were. Here 26% of DA neurons, and 24% of FSI’s were phase-locked to 8-13 Hz, comparable to the latter work.

Here, there were no significant differences between incidence of theta phase-locked neurons during the respective trial periods. However, the failure of results to be significant cannot prove the absence of an effect, in particular, in light of our conservative selection criteria. Park and Moghaddam (2017) found coherence between LFPs in VTA and PFC at 5-15 Hz, and this decreased with greater risk of punishment. Thus, both the latter study and Lowe et al (2021) support a role of these rhythmic oscillations in gating of communication between dopaminergic nuclei and striatum (4 Hz) and with neocortex (10 Hz). However, both Fujisawa and Buzsáki (2011) and Duverci, et al. (2018) found that VTA activity at hippocampal theta frequency was not associated with working memory performance, in contrast with the 4 Hz band. This is consistent with the proposal that the different frequency bands could coordinate communication of complementary types of information (Akam and Kullman, 2014).

### 4.4 Beta and gamma rhythms

Relatively few neurons were phase-locked to these bands of DA nucleus LFP’s, and this likely contributed to the absence of significant differences between incidences between cell types and task events. Fujisawa and Buszáki (2011) found high gamma (30-80 Hz) coherence between PFC and the VTA. The narrow 50 Hz band in the choice arm persisted in both their working memory and control tasks.

## 5 Conclusion and Perspectives

In summary, there is oscillatory activity at several different frequency bands in the dopaminergic VTA and SNc. Of particular interest are the slow (low delta) and 4 Hz bands since their time scale bridges behavioral and neural events. One mechanism for this would be by amplitude modulation of higher frequency oscillations by the phase of slow oscillations, as observed in other systems (Jensen and Colgin, 2007). However, before investigating this, it would be helpful to first determine where the origin of slow oscillation in the dopaminergic nuclei. It would also be interesting to investigate coherence of VTA/SNc oscillations with prefrontal cortex, striatum, and hippocampus as well as other closely associated structures such as lateral habenula for this, and other, frequency bands. Indeed, the VTA and lateral habenula have reciprocal connections, and lateral habenula neuronal activity is phase-locked to the hippocampal theta rhythm (Goutagny et al 2013). Overall, these oscillatory activities would provide a mechanism of potential utility for dopaminergic coordination of brain activity for goal-directed decision-making, learning, and motor control (Williams, et al., 2002).

## 7 Author contributions

JM and SW designed and performed the experiments; JM constructed the experimental apparatus, informatics control and data acquisition systems with support from HN; MV and LV maintained and genotyped the mouse line and guided immunohistochemical processing; FP guided adaptation of the maze and behavioral protocols for training and recording; JM, VO and SIW analyzed data with support from HN, and wrote the manuscript. All authors approved of the manuscript.

## 9 Acknowledgements

Thanks to France Maloumian for help with figures, to Yves Dupraz for constructing the experimental chamber, Estelle Anceaume for microscopy training, Michaël Zugaro for helpful suggestions and informatics support, Dr. Guillaume Dugué and NeuroFabLab for facilitating with 3D printing of headstages, Drs. Karim Benchenane and Liyang Xiang for help setting up the optical stimulation, and Dr. Philippe Faure for advice on recording dopaminergic neurons with chronically implanted octrodes.

## 10 Contribution to the field statement

Rhythmic oscillations of neuronal excitability can coordinate activity within and across brain structures. This is of particular interest for dopaminergic neurons since they impact on learning of both motor sequences and goal-directed behavior. Here we characterize and compare several frequencies of oscillations of local field potentials and single unit activity in brainstem dopaminergic nuclei, highlighting some behavioral correlates. In order to definitively determine whether neurons were dopaminergic, a transgenic mouse line was used, permitting optogenetic identification. In order to characterize functional properties, local field potentials and neurons were recorded in mice training in operant olfactory and visual discrimination tasks. Detection of phase-locking with two methods (Rayleigh test and Pairwise Phase Consistency, PPC) permitted conclusive evidence that oscillatory activity was intrinsic to the dopaminergic nuclei. To avoid errors in tallying responsive neurons, a conservative method excluded putatively repeated recordings of neurons on successive days. Unexpectedly, phase-locking was most common in the slow (awake delta) band. Phase-locking to certain bands occurred more during specific periods in the task. Since dopamine release is implicated in pathologies including Parkinson’s disease, drug and gambling addictions, obsessive-compulsive disorder, etc., future work facilitating or impairing this oscillatory activity could have translational applications.

