## Supplementary material for "Rhythmic oscillations in the midbrain dopaminergic nuclei in mice": Supp Fig 1

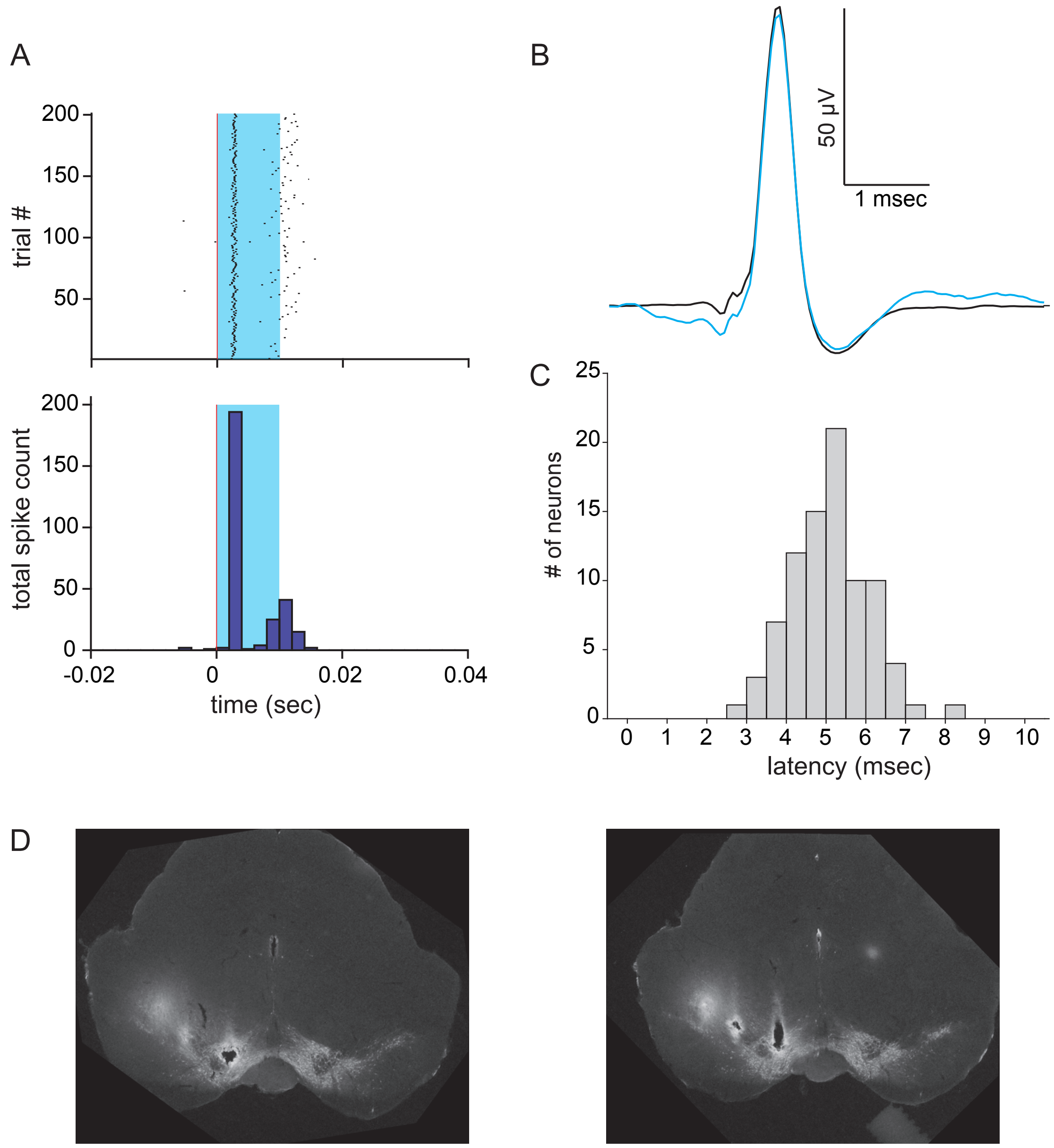


Supp. Fig 1. Recordings in optogenetically identified dopaminergic nuclei in transgenic mice. A) Example of a dopaminergic neuron response to light stimuli (blue shaded period). Spikes appear above, and summary histograms below. B) Waveform of the neuron in A. C) Mean latencies of first spikes for all responsive neurons recorded. D) Example of tyrosine hydroxylase histochemistry of coronal brain sections with electrode tracks and electrolytic marking lesions. The lateral-most electrode did not reach the SNc in this mouse.
