## Supplementary material for "Rhythmic oscillations in the midbrain dopaminergic nuclei in mice": Supp Fig 2

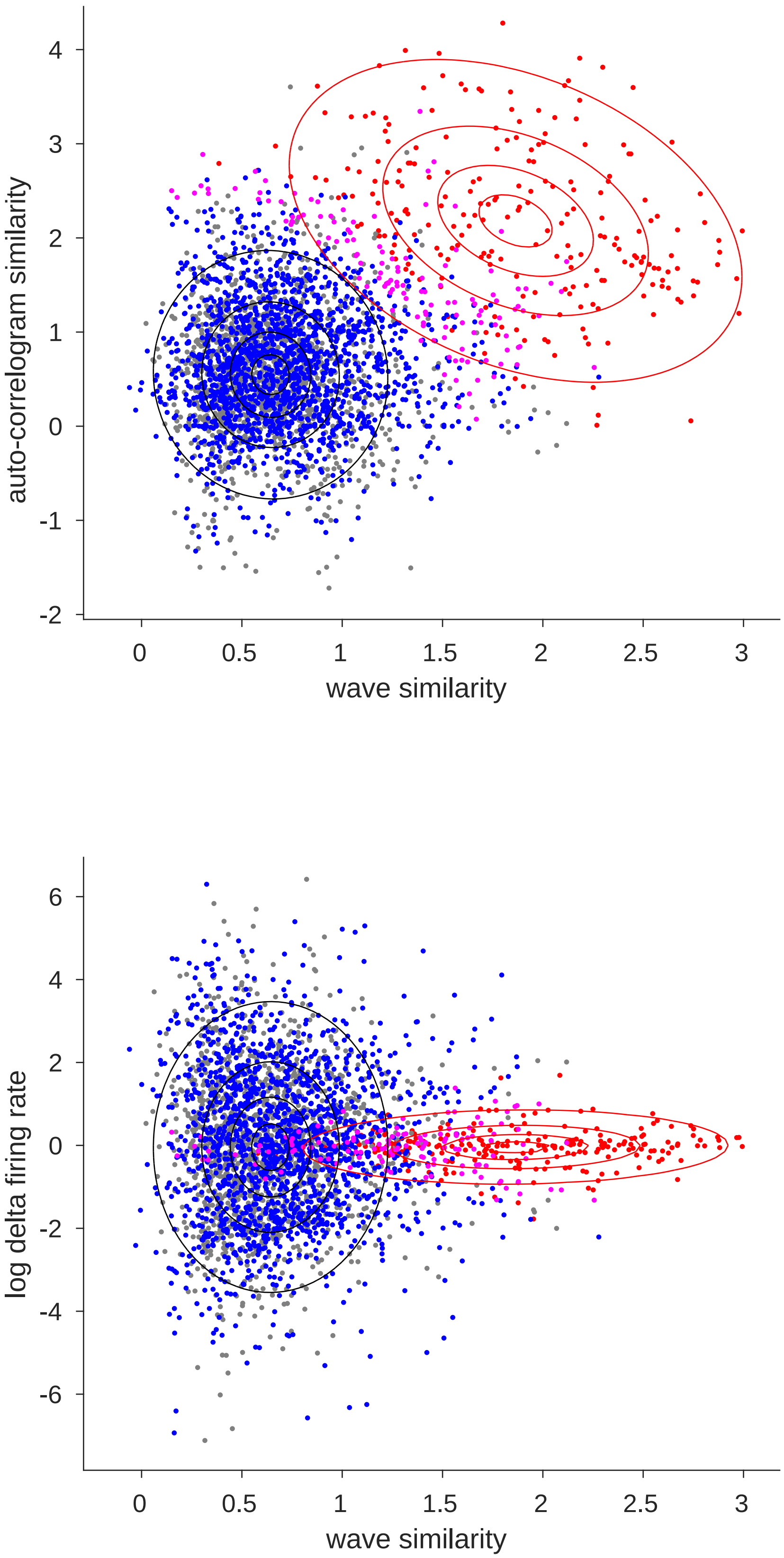


Supp. Fig. 2. Classification of different- and same-neuron pairs recorded in consecutive days. Each dot represents similarity scores of electrophysiological features of a pair of neurons recorded in consecutive days. Dark gray dots are pairs recorded from different octrodes; blue and red dots are pairs recorded from the same octrodes estimated as different- and same-neuron pairs, respectively; pink dots are unknown pairs. The estimated Gaussians of different- (blue) and same-neuron (red) pairs are shown as contour plots. The contours correspond to 25%, 50%, 75%, and 95% of each distribution.
