## Supplementary material for "Rhythmic oscillations in the midbrain dopaminergic nuclei in mice": Supp Fig 3

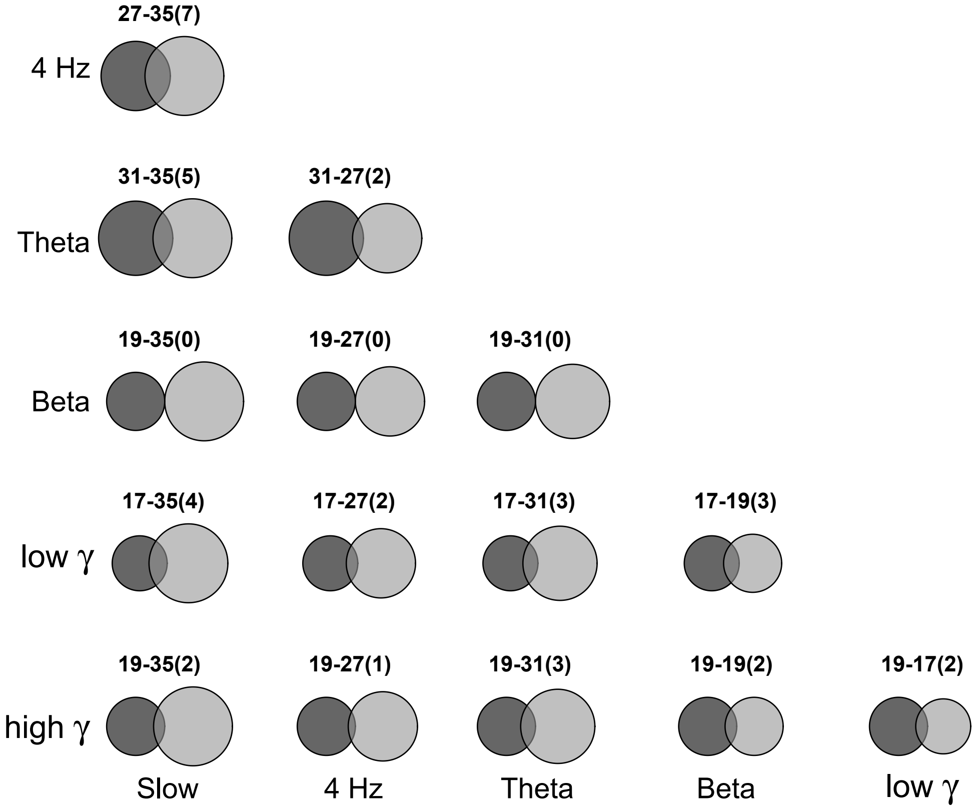


Supp. Fig. 3. Low incidence of phase-locking of the same neuron to two different frequency bands. Venn diagrams indicate the numbers of neurons phase-locked to the bands listed to the left by the area of the black circles, and phase-locked to the bands listed below by the area of the light gray circles. Overlap (dark gray) indicates the neurons phase-locked to both bands. Respective numbers are shown above, with overlap in parentheses.
