## Supplementary material for "Rhythmic oscillations in the midbrain dopaminergic nuclei in mice": Supp Fig 4

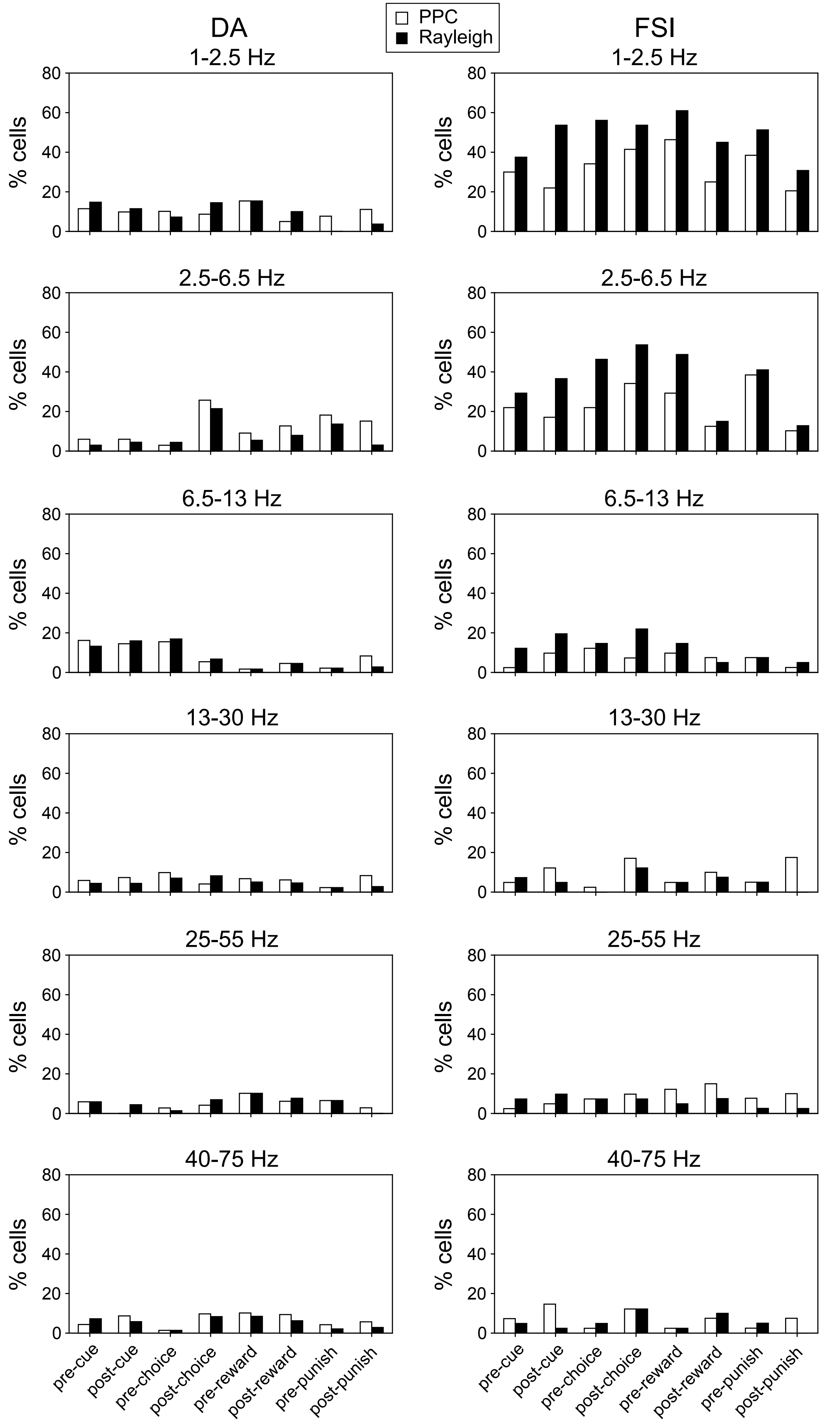


Supp. Fig. 4. Similar profiles in Rayleigh test and PPC for significant phase locking in neurons recorded in the midbrain dopaminergic nuclei during the behavioral task.
